# Strong gamma frequency oscillations in the adolescent prefrontal cortex

**DOI:** 10.1101/2021.08.20.455686

**Authors:** Zhengyang Wang, Balbir Singh, Xin Zhou, Christos Constantinidis

## Abstract

Working memory ability continues to mature into adulthood in both humans and non-human primates. At the single neuron level, adolescent development is characterized by increased prefrontal firing rate in the delay period, but less is known about how coordinated activity between neurons is altered. Local field potentials (LFP) provide a window into the computation carried out by the local network. To address the effects of adolescent development on LFP activity, three male rhesus monkeys were trained to perform an oculomotor delayed response task and tested at both the adolescent and adult stage. Simultaneous single-unit and LFP signals were recorded from areas 8a and 46 of the dorsolateral prefrontal cortex (dlPFC). In both the cue and delay period, power relative to baseline increased in the gamma frequency range (32 - 128 Hz). In the adult stage, high-firing neurons were also more likely to reside at sites with strong gamma power increase from baseline. For both stages, the gamma power increase in the delay was selective for sites with neuron encoding stimulus information in their spiking. Gamma power and neuronal firing did not show stronger temporal correlations. Our results establish gamma power decrease to be a feature of prefrontal cortical maturation.

**Significance Statement:** Gamma-frequency oscillations in extracellular field recordings (such as LFP or EEG) are a marker of normal interactions between excitatory and inhibitory neurons in neural circuits. Abnormally low gamma power during working memory is seen in conditions such as schizophrenia. We sought to examine whether the immature prefrontal cortex similarly exhibits lower power in the gamma frequency range during working memory, in a non-human primate model of adolescence. Contrary to this expectation, the adolescent PFC exhibited stronger gamma power during the maintenance of working memory. Our findings reveal an unknown developmental maturation trajectory of gamma band oscillations and raise the possibility that schizophrenia represent an excessive state of prefrontal maturation.

## Introduction

Human executive functions including working memory continue to mature after the onset of puberty (Fry and Hale, 2000; Gathercole et al., 2004; Davidson et al., 2006; Ullman et al., 2014). This pattern of maturation parallels continued structural changes in the prefrontal cortex: early childhood excitatory synaptogenesis is followed by synaptic pruning in adolescence (Bourgeois et al., 1994; Anderson et al., 1995; Huttenlocher and Dabholkar, 1997), along with decreasing cortical thickness and gray matter volume (Giedd and Rapoport, 2010). In tasks requiring working memory, human imaging studies have also reported distinct changes in prefrontal activity patterns in humans between childhood and adulthood (Luna et al., 2001; Bunge et al., 2002; Klingberg et al., 2002; Kwon et al., 2002; Olesen et al., 2003; Burgund et al., 2006; Olesen et al., 2007). Other markers of neuronal activity, including the distribution of power across different frequency bands of the EEG also changes markedly between the time of adolescence and adulthood (Uhlhaas et al., 2009). Gamma-band oscillations are thought to be driven primarily by excitatory-inhibitory neuronal loops (Buzsaki and Wang, 2012), thus suggesting that strength of gamma oscillations can serve as a marker of synaptic maturation (Uhlhaas et al., 2010). Schizophrenia, a neurodevelopmental disorder with a typical early adulthood onset are characterized by impaired working memory performance (Goldman-Rakic, 1994), abnormal trajectory of prefrontal interneuron maturation (Dienel and Lewis, 2019) as well as decreased power of gamma oscillations (Woo et al., 2010; Uhlhaas and Singer, 2013).

Non-human primates exhibit a strikingly similar pattern of cognitive development and prefrontal maturation (Constantinidis and Luna, 2019) and allow for more direct insights in the nature of neural changes that mediate these phenomena. Single-neuron recordings obtained in non-human primates at different developmental stages have thus revealed increased firing rate specifically during the intervals of memory maintenance (Zhou et al., 2013; Zhou et al., 2014; Zhou et al., 2016b). Continued maturation of inhibitory connections has also been directly documented in the adolescent prefrontal cortex (Gonzalez-Burgos et al., 2015), as have been changes in the effective, intrinsic connectivity between prefrontal neurons between adolescence and adulthood (Zhou et al., 2014). Analysis of neuronal rhythmicity in the adolescent primate prefrontal cortex has not been reported until now. Local filed potentials (LFP), a signal representative of the summation of postsynaptic activity in a small cortical volume (Kajikawa and Schroeder, 2011), provide a mesoscopic measure to link these empirical results. As in human EEG studies, working memory maintenance is generally characterized by elevated gamma-frequency power in the local field potential (Pesaran et al., 2002; Howard et al., 2003; Jensen et al., 2007; Honkanen et al., 2015; Kornblith et al., 2016).

We were thus motivated to determine whether prefrontal cortical maturation between adolescence and adulthood is characterized by increased gamma-band rhythmicity in the local field potential. We analyzed LFP recordings obtained from the same subjects as the transitioned from adolescence to adulthood (Zhou et al., 2016b; Zhou et al., 2016c). Unexpectedly, we found robust gamma band oscillations in the adolescent prefrontal cortex that declined rather than increased during adulthood.

## Methods

### Subjects and data collection

Neurophysiological data used for analysis was collected from male rhesus monkeys performing a visual working memory task and reported in detail previously (Zhou et al., 2013; Zhou et al., 2014; Zhou et al., 2016a; Zhou et al., 2016c). All surgical and animal use procedures were reviewed and approved by the Wake Forest University Institutional Animal Care and Use Committee, in accordance with the U.S. Public Health Service Policy on humane care and use of laboratory animals and the National Research Council’s Guide for the care and use of laboratory animals. Quarterly morphometric and hormonal measures were obtained to determine each animal’s onset of puberty as well as maturity.

The monkeys were trained to perform the ODR task (Fig. 1). This task required the subject to remember the location of a 1° white square stimulus presented for 0.5 s following a 1 s fixation period. The stimulus could appear at one of eight locations arranged on a circle of 10° eccentricity. After a 1.5 s delay period, the fixation point was extinguished, prompting the subject to saccade to the remembered location within a 0.6 s time window to receive a liquid reward. The saccade endpoint had to deviate no more than 5-6°from the center of the stimulus. Correct trials were included for subsequent analysis.

**Figure 1.**
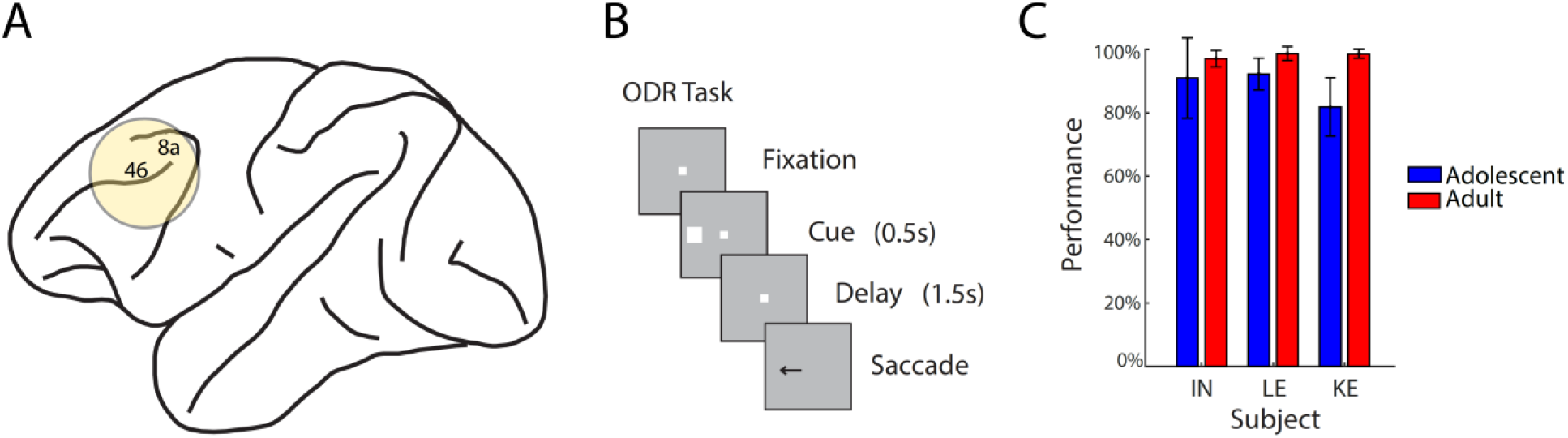
Brain regions recorded, task design and behavior performance. A: Schematic diagram of the monkey brain with locations of neurophysiological recordings in the dorsolateral prefrontal cortex indicated (areas 8a and 46). B: The oculomotor delayed response task required the subject to saccade to a remembered location after a 1.5 second delay. C: Behavior performance of the three subjects used in this study in two stages 1.6 – 2.1 years apart, respectively referred to as the “adolescent” and “adult” stage. Mean percentage of correct trials in ODR task, excluding breaks in fixation, for young: 88%; adult: 98%. Error bars represent standard deviation.

Neural recordings were collected from areas 8a and 46 of the dorsolateral prefrontal cortex. Epoxylite-coated Tungsten electrodes with a diameter of 250 μm and an impedance of 4 MΩ at 1 KHz (FHC Bowdoin, ME) were acutely advanced into the brain, and a reference electrode was attached to the metal recording chamber. Spike waveforms were amplified, band-pass filtered between 0.5 and 8 kHz, and sampled at 40 kHz, while unipolar LFP traces were sampled continuously at 500 Hz. Both signals were digitized and stored through a modular data acquisition system (APM system, FHC, Bowdoin, ME).

### LFP signal processing

LFP recordings were processed using the FieldTrip (https://www.fieldtriptoolbox.org/) and Chronux (http://chronux.org/) toolboxes as well as custom MATLAB code in MATLAB 2018b (MathWorks). LFP signals of individual trials first underwent artifact rejection. Single-trial LFP traces were zero-meaned and then band-pass filtered between 1 and 200 Hz with a zero-phase 5^th^ order Butterworth filter (10^th^ order in effect) and then notch-filtered at 60 Hz with a bandwidth of 0.2 Hz using the *ft_preproc_dftfilter* function. The signals were then standardized by their standard deviation as estimated by the median absolute deviation. The spectrogram of single traces between 2 and 128 Hz were computed by the *mtspecgramc* function with 6 tapers of 500 ms time windows. LFP power in the alpha, beta, gamma and high-gamma bands were defined to be the sum of power in the spectrogram between the frequencies of 8 – 16 Hz, 16 – 32 Hz, 32 – 64 HZ and 64 – 128 Hz respectively. We used the last 500 ms of the fixation period as the baseline period. Relative band power was calculated by dividing the sum of power within each band at each time point by the average of the sum in the baseline period. The values in the relative spectrograms (Fig. 2) were calculated separately at each frequency (at a 2-Hz resolution). To identify recording sites where LFP power was modulated in a specific frequency band, a two-tailed paired-sample t-test was carried out on the band power from all trials at each recording site averaged during the delay period. Sites with p values below 0.05 was defined to be modulated in the corresponding frequency band. To find spatially selective LFP recording sites, 1-way ANOVA test was carried out on the band power of each recording site averaged during the delay period in response to different stimulus location. LFP recording sites with p values below 0.05 were defined to be selective for stimulus location in the corresponding power band.

**Figure 2.**
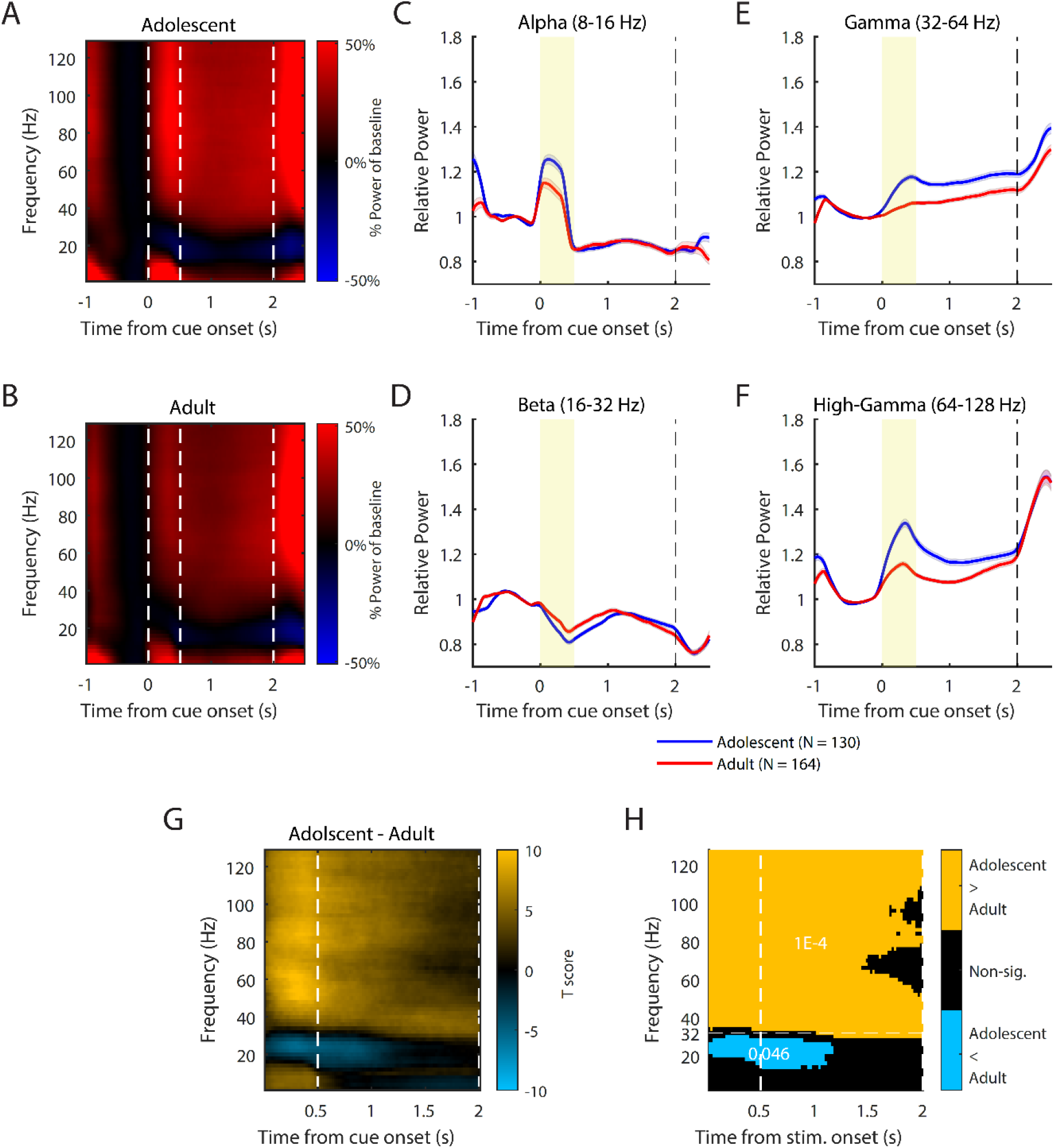
dlPFC LFP power changes during adolescent development. A, B: Population average across electrodes and sessions of moment-by-moment LFP spectral power in proportion to that in the last 500 milliseconds prior to cue onset for LFP recording sites at the adolescent (N = 130) and adult (N = 164) stage. C-F: Population averages of LFP power evolutions aligned to cue onset in the frequency bands between 8-16, 16-32, 32-64 and 64-128 Hz at sites recorded in the adolescent (N = 130) and adult (N = 164) stage. G: Uncorrected 2-sample t-test t values comparing the moment-by-moment LFP spectral power in proportion to that in the last 500 milliseconds prior to cue onset between the adolescent (N = 130) and adult (N = 164) stage. Positive values correspond to larger adolescent power. H: Significant clusters found in the permutation test based on t values in A. The number in each cluster represents the bootstrapped p value.

### Spike processing

Recorded spike waveforms were sorted into separate units using an automated cluster analysis method based on the KlustaKwik algorithm. Trial-averaged PSTH was computed by convolving the spiking events with a 50-ms boxcar kernel at 20-ms steps apart. The evoked PSTH was calculated by subtracting the average firing rate in the 1-second fixation period. Single-trial PSTH for computing moment-by-moment percentage of explained variance (PEV) used a 250-ms box kernel instead. Neurons were identified to be responsive to the task, if their activity increased significantly during any task epoch relative to the baseline fixation period, evaluated at the 0.05 significance level, as we have described previously (Zhou et al., 2016c). To identify neurons selective for spatial location during the delay period of the task, a 1-way ANOVA test was carried out on the firing rates of each neuron averaged during the entire delay period. Selective neurons were defined to be the ones with p values below 0.05. The bias-corrected PEV *ω*^2^ was computed in equation (1) at steps of 20 ms on the single-trial firing rate of each neuron, where MSE is the mean squared error within stimulus locations, df the degree of freedom, SS_Between_ the sum of squares between stimulus locations and SS_total_ the total variance.

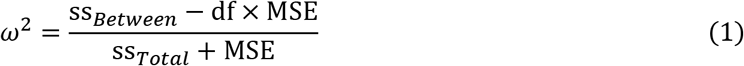

### Cluster-based permutation test

A permutation test with 10,000 iterations was carried out to compare the relative spectrogram between the adolescent and the adult using the FieldTrip function *ft_freqstatistics.* two-tailed independent-sample t-tests were first run for each time-frequency combination to generate uncorrected p values before connected regions of all positive or negative t statistics with p values smaller than 0.05 were identified (clusters). The sum of all t statistics within each cluster were calculated. The labels for developmental stage were then randomly shuffled 10,000 times where each time the largest cluster sum of t values was recorded. The original cluster sum of t values was then compared against the bootstrapped shuffled distribution to generate a one-tailed p value.

### Correlation analysis

The neuron-by-neuron Spearman’s correlation between neuronal firing rate/PEV and LFP power was computed by first averaging the signals within each neuron or recording site and within the delay period and then matching the signals that were recorded from the same electrode in each session. The correlation coefficients were compared across the adolescent and the adult using Fisher’s Z transform.

Temporal correlation was calculated for the time-series signals in the delay period of each trial. The spiking events were convolved with a 50-ms boxcar kernel. For each neuron, the trials for each stimulus location were concatenated and the correlation was computed between the convolved spiking and the LFP power changes from baseline at each frequency. Only results from the stimulus location that resulted in the strongest evoked delay period firing rate for each neuron were further considered. When examined on a frequency-by-frequency basis, all p values were controlled for using the Bonferroni-Holm procedure (see the Statistics section for details).

### Statistics

All statistical tests were conducted using MATLAB 2018b. An alpha level of 0.05 was adopted for all tests. The family-wise error rate (FWER) was controlled for using the Bonferroni correction when multiple comparisons were conducted in multiple canonical frequency bands. For *n* comparisons, the Bonferroni correction sets the critical p value to *α/n*. Alternatively, the Bonferroni-Holm procedure was carried out when multiple comparisons were conducted in a frequency-by-frequency fashion. Briefly, the Bonferroni-Holm procedure sorts all p values from lowest to highest, then compares in order from the first to the *k*th p value to a critical level of *α/(n* +1 – *k*). A comparison is considered significant if the current p value is smaller than the critical level, otherwise the current and all following comparisons are deemed not significant, and the procedure stops.

## Results

Data were analyzed from three male Rhesus monkeys (*Macaca mulatta*), trained to perform the oculomotor delayed response (ODR) task (Fig. 1B). The task tested the animals’ visual working memory ability by requiring the subject, after a delay of 1.5 second, to make a saccade to a remembered stimulus location that was presented for 0.5 seconds. The animals were tested at two stages of development: the ‘adolescent’ stage after puberty onset (approximately age 4.5 years) and the ‘adult’ stage once development had completed (approximately age 6.5 years). Young monkeys generally achieved a lower performance in the task than adult ones (Fig. 1C, 88% vs. 98% correct performance for young and adult, respectively, not considering aborted trials), as we have reported previously (Zhou et al., 2016c). During performance of the task, spiking and LFP activity was recorded from area 8a and 46 (Fig. 1A). We analyzed LFP activity from electrodes where single unit activity was also identified. The resulting dataset consisted of signals from 298 neurons and 130 LFP sites in the adolescent stage along with 392 neurons and 164 LFP sites in the adult stage.

### Adolescent dlPFC shows a stronger pattern of LFP power modulation in the ODR task

We calculated LFP spectral power at each epoch of the ODR task, relative to baseline. For each recording session from one electrode, the baseline period for LFP analysis was defined to be the last 500 milliseconds of the fixation period in order to avoid including the transient power changes following fixation onset. The relative power of the recording was then calculated as the spectrogram change from the baseline average at each respective frequency. Results from different sessions and electrodes were averaged together for the adolescent (Fig. 2A) and the adult stage (Fig. 2B) respectively. In order to describe in detail its temporal evolution throughout the different epochs of the task, LFP power modulation was evaluated in four different frequency bands: alpha (8 – 16 Hz), beta (16 – 32 Hz), gamma (32 – 64 Hz) and high-gamma (64 – 128 Hz). The resulting relative power spectrogram showed similar patterns of modulations in each task epoch at the adolescent and the adult stage. Specifically, gamma (Fig. 2E) and high-gamma (Fig. 2F) power were elevated relative to baseline during both the cue presentation and the delay period. Beta power (Fig. 2D) generally showed modulations in the opposite direction as gamma, showing decreases in both the cue and delay periods. Alpha band power (Fig. 2C) had an initial increase during cue presentation and then quickly dropped below baseline in the delay period.

Contrary to our initial hypothesis, gamma-band and high-gamma band LFP power was more strongly elevated during the delay interval of the task in the adolescent than the adult stage (Fig. 2E-F). Averaged across the entire duration of the delay period, the difference was highly significant for the gamma (two-tailed t-test t(292) = 5.09, p = 6.4E-7) and high-gamma range (two-tailed t-test t(292) = 4.86, p = 1.9E-6). This difference was specific for the gamma and high-gamma frequency ranges. No significant difference was detected for the alpha or beta power (two-tailed t-test t(292) = −0.13 and −0.68 respectively; p = 0.89 and 0.50 respectively). In addition, as previously reported, adult dlPFC neurons featured higher firing rates, both in terms of absolute firing rate and as an increase over the baseline. This was true for the subset of neurons (Fig. 4A, C) recorded in electrode penetrations where these LFPs were recorded. As a result, the difference in gamma and high-gamma power between the two stages cannot be accounted for by systematic differences in firing rate and in fact moved in the opposite direction than would be expected based on firing rate alone.

Such differences between stages in spectral power changes during the cue presentation and memory delay were further confirmed using a cluster-based permutation test (Fig. 2G, H) at every 2 Hz between 2 and 128 Hz. Consistent with findings in bandpass power, one strong cluster was identified: LFP power modulation above around 30 Hz was significantly higher (p = 1.0E-4) for the adolescent than the adult throughout the cue presentation and memory delay epochs. One other marginally significant (p = 0.0457) cluster was found suggesting stronger negative modulation for the adolescent below 30 Hz in an epoch immediately following cue presentation.

In order to rule out potential biased sampling of recording sites with heterogeneous LFP power properties, we defined a site to be LFP power-modulated if its delay period power in a certain frequency band differed significantly relative to baseline (evaluated with a paired t-test, at the α=0.05 significance level). Most adolescent and adult sites were gamma power modulated, but more so adolescent (adolescent: 104/130; adult: 97/164, Fisher’s exact test, p = 1.4E-4). Similar percentage of adolescent and adult sites were power-modulated in high-gamma (adolescent: 117/130; adult: 122/164, Fisher’s exact test, p = 8.1E-4). We repeated the comparison of gamma power between the young and adult stages, including in the analysis exclusively the power-modulated sites. A higher power in the adolescent than in the adult stage was still present in the gamma (Fig. 3A, two-tailed t-test t(199) = 3.07, p = 2.5E-3) and high-gamma frequency band (Fig. 3B, two-tailed t-test t(237) = 3.31, p = 1.1E-3). The observed differences in gamma and high-gamma delay power were thus unlikely due to unequal sampling of modulated vs. non-modulated sites.

**Figure 3.**
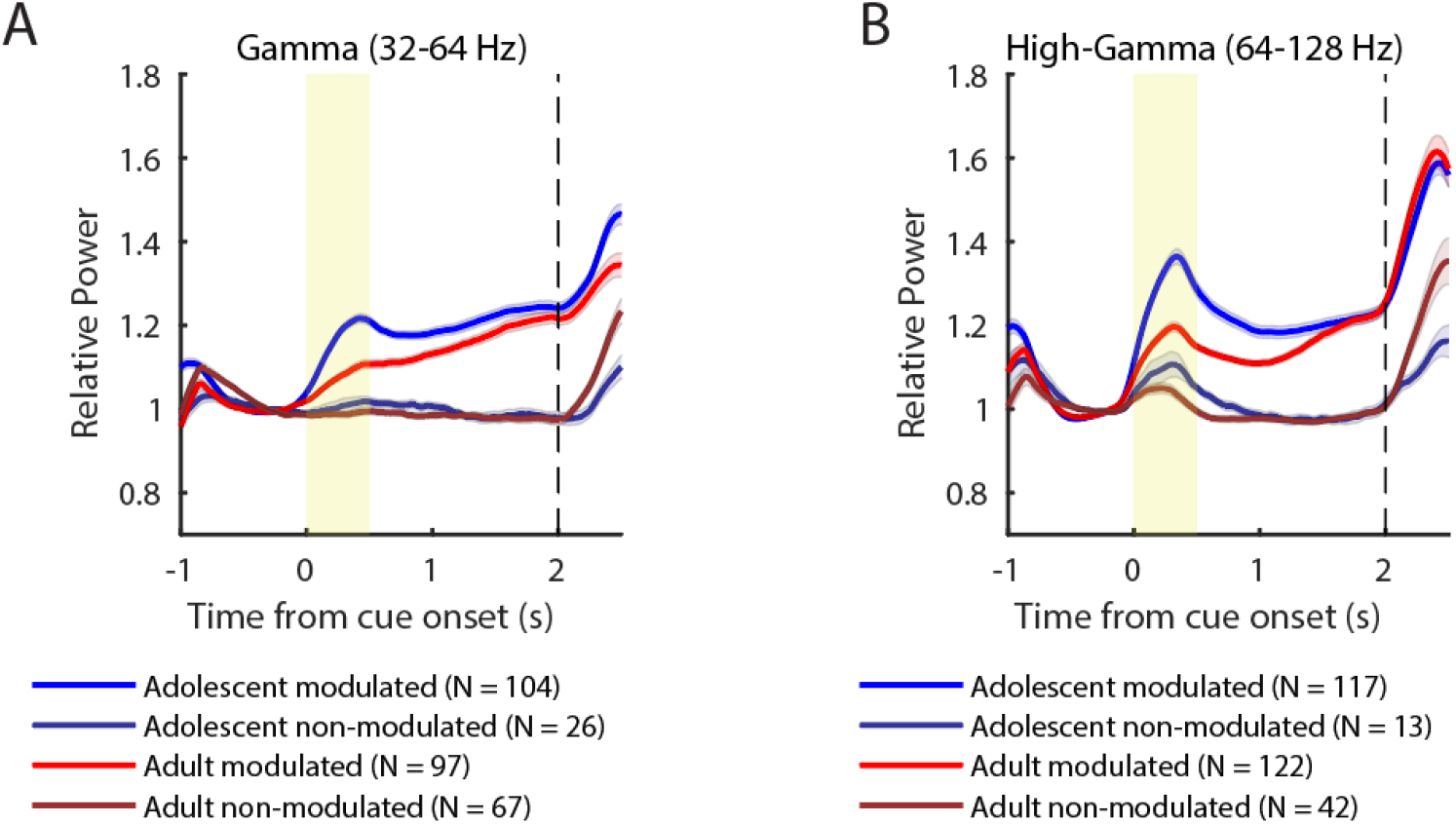
Heterogeneity in gamma and high-gamma LFP power changes across recording sites. A, B: LFP band power evolutions of sites grouped according to the level of changes (significant/non-significant) in gamma (adolescent 104/26, adult 97/67) or high-gamma (adolescent 117/13, adult 122/42) power respectively during the working memory delay. Shaded areas represent SEM.

**Figure 4.**
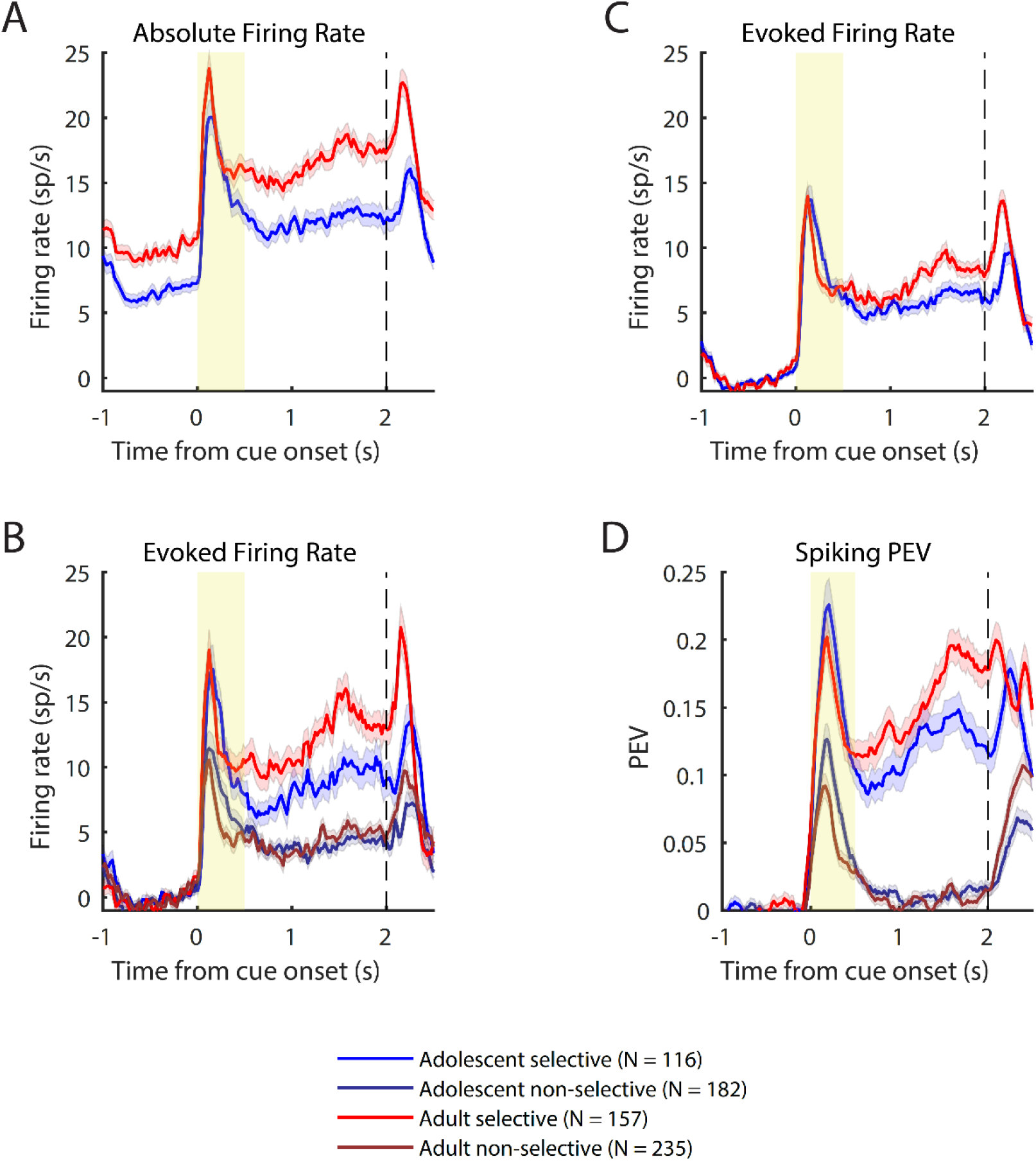
dlPFC neuronal spiking and information encoding in the ODR task. A, C: Population averages of neuronal raw firing rate and evoked firing rate by subtracting the average in the 1-second fixation period for neurons recorded in the adolescent (N = 298) and adult (N = 392) stages. B, D: Average evoked firing rate and proportion of its explained variance (PEV) by cue location after dividing the neurons according to differential delay period firing (selective/non-selective) in the adolescent (116/182) and adult (157/235) stage.

LFP power in the delay period is often tuned for the spatial location of the stimulus (Pesaran et al., 2002). We therefore wished to test whether the higher adolescent gamma power was only evident in sites selective for the stimulus location or in sites that were not modulated by the stimulus. We defined spatially-selective LFP sites as those in which delay period power differed significantly depending on the location of the stimulus (evaluated with 1-way ANOVA at the α=0.05 significance level). Similar percentages of adolescent and adult LFP sites exhibited spatially-selective gamma power (adolescent: 31/130; adult: 28/164) and high-gamma (adolescent: 41/130; adult: 52/164). In the spatially-selective sites, gamma and high-gamma delay-period LFP power was higher overall in the adolescent than the adult stage, but the difference did not reach significance for the gamma range (two-tailed t-test t(57) = 0.78, p = 0.44); it only did for high-gamma power (two-tailed t-test t(91) = 3.55, p = 6.1E-4). Among non-spatially-selective sites, delay-period LFP power was significantly higher in the adolescent stage for both the gamma (two-tailed t-test t(233) = 5.31, p = 2.6E-7) and high-gamma frequency band (two-tailed t-test t(199) = 3.89, p = 1.4E-4).

### Adolescent neurons’ delay period firing correlates less with gamma

Gamma oscillations have been suggested to reflect the underlying synchronization of local networks giving rise to delay period activity as well as differential encoding of stimulus information in working memory (Roux and Uhlhaas, 2014). Given that we have previously reported higher firing rate in the delay period during the adult stage (Zhou et al., 2016c), it seemed curious that delay period gamma power and firing rate saw changes in opposite directions during adolescent development. Therefore, we tested for potential changes in correlation at the neuronal population level between gamma power modulation and neuronal spiking. A neuron’s preferred stimulus location was defined as the one that evoked the highest average firing rate in the cue and delay periods combined. The correlation between neuronal firing and LFP power was overall higher in the adult and emerged amongst a broader band of frequencies (Fig. 5A). In the adult stage, the neurons’ evoked delay period firing rates to the preferred stimulus location weakly but significantly correlated with gamma (Spearman’s rho = 0.326, p = 4.0E-11) and high-gamma power modulations in LFP signals recorded at the corresponding sites (Spearman’s rho = 0.513, p = 1.2E-27). The strength of such correlations was weaker in the adolescent stage for the gamma (Spearman’s rho = 0.170, p = 0.003) and high-gamma (Spearman’s rho = 0.360, p = 1.6E-10) bands respectively. The adult correlation between delay period firing rate and high-gamma band LFP was significantly higher than the young one (Fisher’s Z-transform, gamma: p = 0.031 > 0.05/2; high-gamma: p = 0.014 < 0.05/2 under Bonferroni correction).

**Figure 5.**
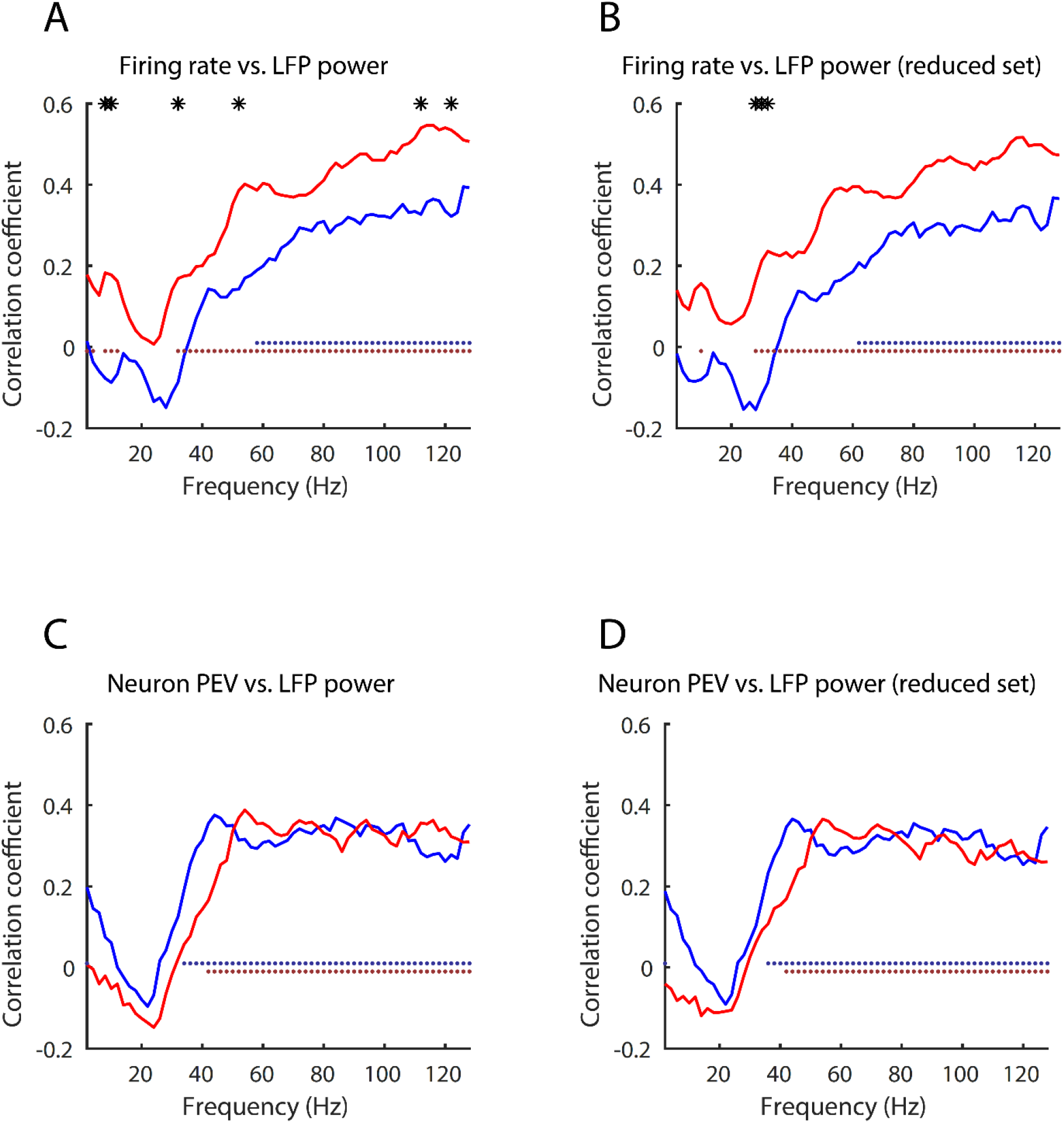
Neuron-by-neuron correlations between spiking activity and LFP power. A: Spearman’s correlation coefficient between neuronal firing rate and LFP power for signals recorded from the same electrode. C: Spearman’s correlation coefficient between neuronal PEV and LFP power for signals recorded from the same electrode. B and D: Same as A and C but for a firing rate-matched subset of neurons. Solid dots indicate significant correlations (Bonferroni-Holm-corrected type I error rate < 0.05). Asterisks stand for significant differences between the adolescent and the adult (Fisher’s Z transform, Bonferroni-Holm-corrected type I error rate < 0.05).

In order to discount the potential contribution of firing rate differences (adolescent: 5.79 ± 7.04 vs. adult: 7.44 ± 8.28 spikes per second) to gamma power through spectral leakage, we assembled an evoked firing rate-matched subset of data by removing the top 5.6% (22/392) of adult neurons and bottom 5.0% (15/298) of adolescent neurons in terms of evoked firing rate in the delay period to the preferred stimulus location. These remaining adolescent neurons had an average evoked firing rate of 6.16 ± 7.04 spikes per second, matching the 6.01 ± 5.49 spikes per second of the remaining adult neurons (two-tailed t-test, t(651) = 0.32, p = 0.75). The reduced dataset’s delay evoked firing showed a very similar pattern of correlations (Fig. 5B) as the full dataset between gamma (adolescent: Spearman’s rho = 0.160, p = 0.007; adult: Spearman’s rho = 0.335, p = 4.0E-11) and high-gamma power modulations (adolescent: Spearman’s rho = 0.339, p = 4.9E-9; adult: Spearman’s rho = 0.490, p = 1.0E-23). After matching for evoked neuronal firing rates, the adult stage still showed significantly stronger correlations than the adolescent one (Fisher’s Z-transform, gamma: p = 0. 018 < 0.05/2; high-gamma: p = 0.021 < 0.05/2 under Bonferroni correction). Furthermore, these differences in correlation cannot be explained by the mismatch of the two stages’ gamma power either, for the adolescents’ higher gamma power average should lead to higher correlation coefficients if the underlying true distribution of correlations were the same between the two stages. The results of this analysis suggest that network activity may be constraining unit responses more in the adult than the adolescent stage.

### Delay period neuronal information encoding is associated with greater gamma power

In analogy to spatially-selective LFP sites, we identified spatially-selective single neurons if their average evoked firing rates in the delay period were significantly different across stimulus locations and ‘non-selective’ otherwise (see Methods). This resulted in 39% (116 out of 298) of adolescent neurons and 40% (157 out of 392) of adult neurons being categorized to be spatially-selective. The ratio of selective to non-selective neurons did not differ significantly between the 2 stages (Fisher’s exact test, p = 0.81). Amongst the selective neurons, those in the adult had higher evoked firing rate in the delay period to their preferred stimulus locations (Fig. 4B; two-tailed t-test, t(271) = 2.98, p = 0.003) as well as higher PEV (Fig. 4D; two-tailed t-test, t(271) = 2.71, p = 0.007). We used a 2-way ANOVA to compare delay-period LFP power in sites where selective and non-selective neurons were recorded. LFP delay period gamma (Fig. 6C) and high-gamma (Fig. 6D) power was higher at sites where selective neurons were recorded, evidenced by a significant main effect of selective-neuron site (gamma: F(1, 290) = 24.78, p = 1.1E-6; high-gamma F(1, 290) = 25.04, p =9.8E-7). There was also a significant main effect of developmental stage (gamma: F(1, 290) = 24.85, p = 1.1E-6; high-gamma: F(1, 290) = 22.42, p = 3.4E-6), but no interaction between the selective-neuron site and developmental stage (gamma: F(1, 290) = 3.30, p = 0.07; high-gamma: F(1, 290) = 0.16, p = 0.69).

**Figure 6.**
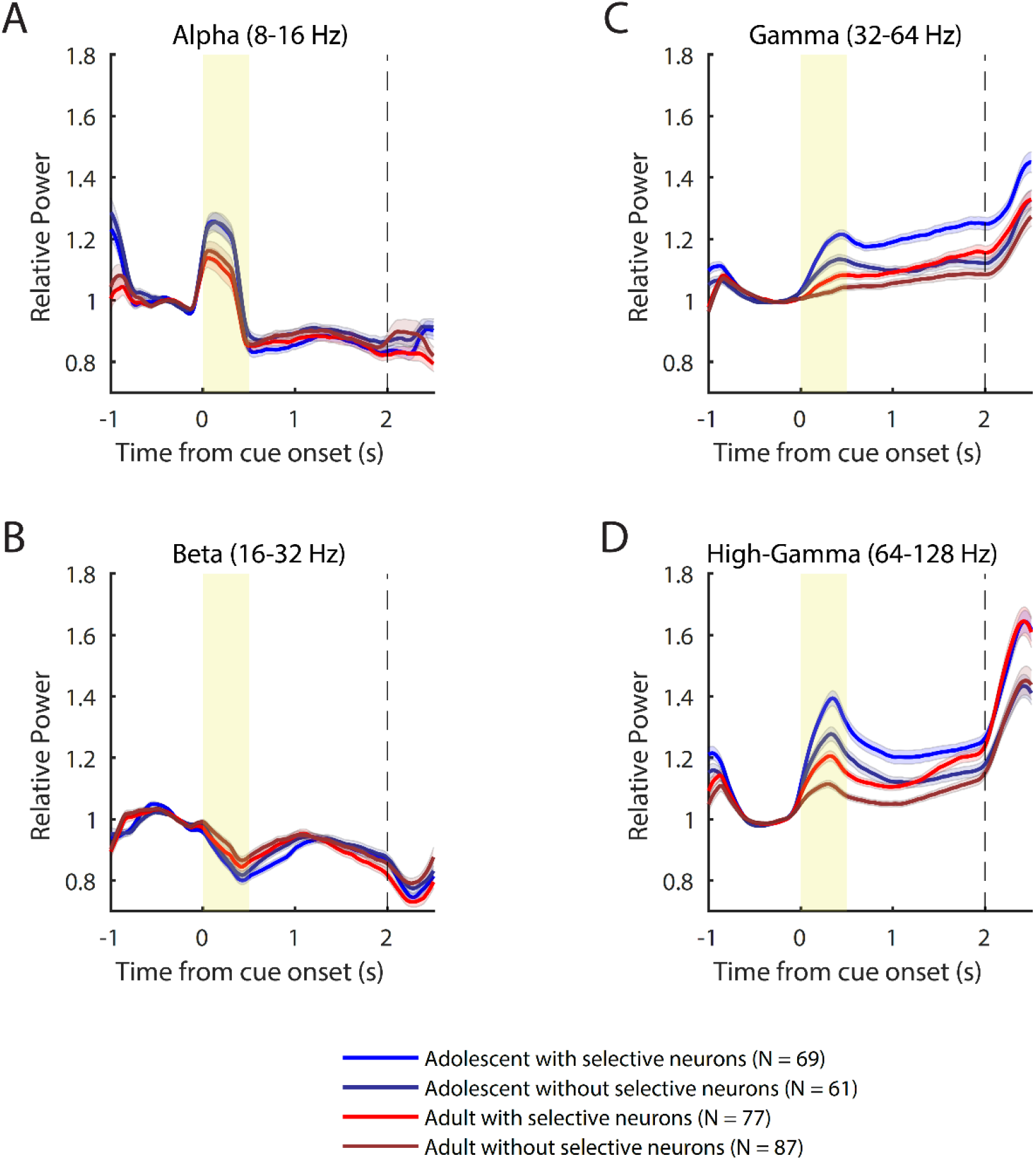
LPF power modulation as a function neuronal information encoding. LFP band power evolutions of sites grouped according to whether spatially-selective neurons were recorded at the corresponding sites in the adolescent (69/61) and adult (77/87) stage.

Such systemic effect could have been underlain by preference of selective neurons for gamma-modulation. Indeed, selective neurons were more likely to be found at gamma-modulated recording sites compared to non-selective neurons in both the adolescent (selective neurons: 95.7%, N = 116; non-selective neurons: 85.7%, N = 182; Fisher’s exact test, p = 0.006) and the adult (selective neurons: 74.5%, N = 157; non-selective neurons: 58.3%, N = 235; Fisher’s exact test, p = 0.001) stage.

A frequency-by-frequency exanimation of the correlation between LFP power and the PEV of neuronal discharges (Fig. 5C) revealed that for both the adolescent and adult neurons, there was significant correlation between neuronal information and LFP power in the gamma and high-gamma frequency range (Bonferroni-Holm-corrected type I error rate < 0.05). Similar to the findings above based on band-passed LFP power, no significant difference was found between the adolescent and the adult at any frequencies (Fisher’s Z transform, Bonferroni-Holm-corrected), indicating the association between gamma power and neuronal information encoding remained highly similar in both developmental stages. For the reduced dataset with matched firing rates for the adolescent and adult neurons, the distribution of correlations was still highly consistent between the two stages (Fig. 5D).

### No strong temporal correlation between LFP power modulations and neuronal spiking

LFP oscillations in the dlPFC have been previously suggested to control neuronal spiking and thus information encoding in a moment-by-moment fashion through their transient temporal dynamics (Lundqvist et al., 2016; Lundqvist et al., 2018). In order to test at the single-trial level, how much the delay period moment-by-moment change in gamma and high gamma power reflected neuronal spiking, we extracted, for each neuron’s preferred stimulus trials, their firing rate (convolved with a 50-ms boxcar kernel) and band power traces. On single trials, gamma power often exhibited short-living peaks surrounded by decreases in power. When single trials from all recording sites were pooled according to developmental stage, gamma power exhibited peak time points throughout the delay period (Fig. 7A-B). Both the adolescent and adult gamma power peak times followed a similar distribution (two-sample Kolmogorov-Smirnov test D(6185) = 0.03, p = 0.131). The trial-by-trial spiking of the matching neurons on the other hand, showed no evidence of activity clustered at these gamma peaks and thus bore little temporal structure similarity (Fig. 7C-D). This was to be expected according to previous accounts of prefrontal neurons exhibiting higher temporal irregularity in the mnemonic delay period with the majority of whose behavior mimicking a Poisson process (Compte et al., 2003). Consequently, the gamma power showed little to no correlation on average with those of neuronal firing at the corresponding recording sites in either the adolescent (Spearman’s correlation, mean rho = 0.045, N = 2792) or adult stage (Spearman’s correlation, mean rho = 0.027, N = 3393). Shuffling the trial pairings between the gamma power and firing rate for 100,000 iterations within in each age group (adolescent 95% confidence interval: −0.004 to 0.011; adult 95% confidence interval: −0.006 to 0.011) showed that the observed mean correlations were significantly higher than chance.

**Figure 7.**
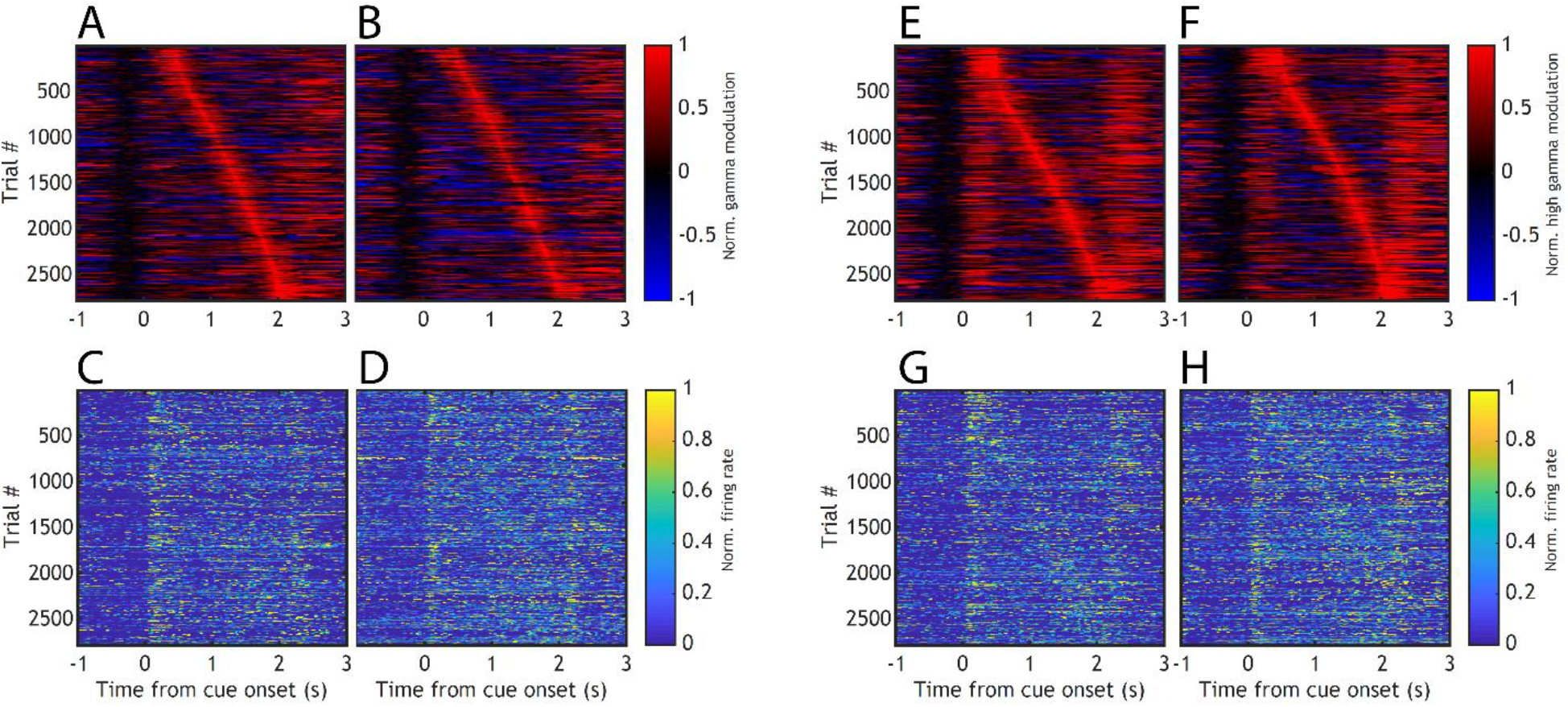
Trial-by-trial temporal evolution of neuronal firing and matched LFP power. A: Adolescent single-trial gamma band power arranged according to peak timing in the delay period (N = 2792). B: Same as A for the adult LFP (N = 3393). C, D: Normalized neuronal firing rate (convolved with a 50-millisecond boxcar kernel) corresponding to the same trials in A and B. E-H: Same as A-D for high gamma power.

Similar analysis was carried out on the temporal profile of high-gamma power (Fig. 7E-H). Interesting, the distribution of peaking time differed significant between the two stages (two-sample Kolmogorov-Smirnov test D(6185) = 0.08, p = 1E-9), with the adolescent high-gamma power peaking earlier overall in the mnemonic delay. Furthermore, there was a weak temporal correlation between high-gamma power and neuronal firing at both the adolescent (Spearman’s correlation, mean rho = 0.111, N = 2792) and adult stage (Spearman’s correlation, mean rho = 0.127, N = 3393). Such correlations were abolished once the trial pairings were shuffled 100,000 times for either the adolescent (95% confidence interval: −0.007 to 0.010) or adult trials ((95% confidence interval: 0 to 0.015).

While the correlation between high-gamma power and firing rate is a common observation (Ray and Maunsell, 2011), the absolute level of correlation observed in this dataset was rather low. Furthermore, spectral leakage from spiking might be responsible for part of this correlation, which would further reduce the magnitude of any true correlations. A frequency-by-frequency inspection of temporal correlations between neuronal firing and LFP powers showed that the correlation coefficient increased with frequencies for both the adolescent and the adult, further suggesting the contribution of spectral leakage from spikes at higher frequencies (Fig. 8). The positive correlations became significantly different from 0 starting at 36 Hz and 42 Hz for the adolescent and the adult respectively. It is also worth noting that, opposite to the developmental effect on neuron-by-neuron correlations, the temporal correlation between LFP power and spiking was higher for the adolescent than the adult, mostly in the high-gamma range. This is further evidence that the stronger association between gamma power and high-firing neurons in the adult was not due to spectral leakage into the LFP directly from spiking activity. In summary, gamma power and neuronal spiking did not follow closely the same temporal profile from trial to trial. The relationship between the temporal dynamics of neuronal stimulus encoding and moment-by-moment LFP power modulation in the delay period showed a wide distribution and were not governed by a single rule across neurons. The adolescent and adult stages only seemed to differ in the distribution of the peak timing of high-gamma power from trial to trial.

**Figure 8.**
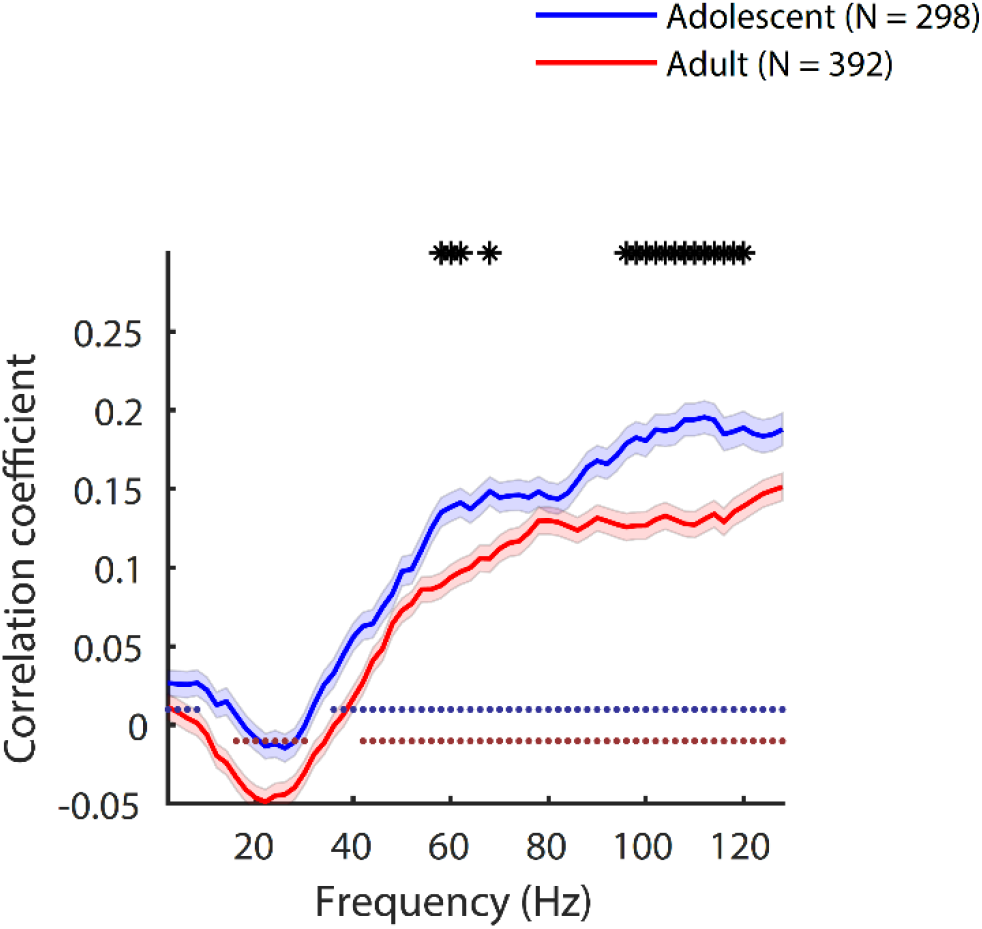
Average temporal correlation of LFP power with neuronal spiking. Solid lines are average Spearman’s correlation coefficient for the adolescent and adult stages respectively. Shaded area represent SEM. Solid dots indicate average correlations significantly different from zero (signed-rank test, Bonferroni-Holm-corrected type I error rate < 0.05). Asterisks stand for significant differences between the adolescent and the adult (Wilcoxon’s rank-sum test, Bonferroni-Holm-corrected type I error rate rate < 0.05).

## Discussion

Here, we analyzed the spiking and LFP signals from the dlPFC of monkeys at the adolescent and adult stage respectively, while they performed a visual working memory task. Previous studies have documented that the adult dlPFC is characterized by increased delay period firing rate (Zhou et al., 2016c), yet we now saw an overall decrease of LFP gamma-band power in the delay period of the task in adult monkeys. A similar change was present for high-gamma power, while alpha and beta band power remained relatively stable during adolescent development. This gamma-power decrease was evident when studying exclusively recordings from sites that exhibited significant gamma power increase in the delay period, suggesting that the findings were not due to any unbalanced sampling of potentially heterogeneous populations of local circuits in the dlPFC. In both the adolescent and adult dlPFC, gamma power was higher at sites where neuronal spiking contained significant stimulus information in the delay period. Interestingly, this separation was largely not interacting with between-stage differences (Fig. 4, 5). These findings, along with the observation that the two stages had almost identical percentages of selective neurons, indicates that peripubertal development of the primate dlPFC does not abolish the already established computational principles in the local circuitry: stronger gamma oscillations would emerge in a local network of neurons actively maintaining task-relevant stimulus information. Rather, the adult PFC differed from the adolescent by having high-firing, information-encoding neurons more heavily distributed at sites with stronger gamma power. This finding, along with the observation that the increase in mature delay period firing rate was biased for selective neurons (Fig. 4) suggests a potential refinement strategy in adulthood: compared to the immature dlPFC, the adult prefrontal cortex more efficiently recruits a task-relevant subset of local networks, engaging them in reverberatory dynamics that are more localized and less diffuse so as to result in lower magnitude of gamma power in population measures such as the LFP. The universal differences seen across all populations of the recording sites are likely to be attributed to overall changes in synaptic properties including synaptogenesis and receptor expression (Lewis et al., 2004; Hoftman and Lewis, 2011; Gonzalez-Burgos et al., 2015), leading to a shift in the excitation-inhibition balance and consequently the level of neural synchronization (Compte et al., 2000).

### Origins of LFP gamma power

The temporal relationship between neuronal spiking and gamma power is complex. During stimulus presentation, the highly correlated bottom-up inputs can serve to synchronize population neuronal spiking and phases of synchronized excitation by pyramidal neurons followed by inhibition by interneurons can thus produce oscillations in the field potentials (Fries, 2009). However, LFP gamma power can also emerge in the delay period, when no sensory input is present, and the postsynaptic potentials are dominated by the local recurrent connections (Pesaran et al., 2002). As modelling findings have suggested, high-frequency oscillations readily emerge in a strongly recurrent network in which single neuron firing is highly irregular, without exhibiting synchronization amongst a sparsely sampled subset of neurons (Wang, 2010). Previous recordings have described dlPFC firing during the delay to be close to Poisson-like (Compte et al., 2003) and Fano factor values above 1 for the adolescent and adult neurons alike (Zhou et al., 2016c). The spatial scope of LFP signals dictates that they reflect the summation of activity from a wide range of neurons (Kajikawa and Schroeder, 2011). Given the highly irregular nature of dlPFC neuronal firing, it is reasonable that LFP would not be highly predictive of activity recorded from a small subset of single neurons from the local population of unsynchronized firing and various stimulus tuning. This was also supported by the fact that a smaller percentage of LFP sites showed significant spatial tuning compared to that of neuronal spiking.

Local-circuit differences have been described between adolescent and adult monkeys that could explain changes in persistent discharges and gamma-band LFP oscillations between developmental stages (Li et al., 2020). Zero-lag spiking synchronization based on cross-correlation analysis of nearby neurons (recorded at distances between 0.5 – 1 mm from each other) is markedly lower in adolescent than in adult monkeys. This difference is primarily the effect of changes in inhibitory interactions, the net efficacy of which declines in adulthood (Zhou et al., 2014). Anatomical evidence, in turn, implicates decreases in the connectivity strength of pyramidal neurons onto interneurons, which lessens the net output of inhibitory connections as the prefrontal cortex matures (Gonzalez-Burgos et al., 2015). This pruning of synaptic interactions implies less synaptic drive overall, which could explain our current finding of decreased rather than increased gamma-band power in the LFP.

### Cognitive processes related to gamma power maturation

Results from human EEG/MEG literature have indicated systematic changes in rhythmicity between developmental stages, generally suggestive of higher gamma power in adulthood. However, closer examination of these studies reveals that existence and direction of change in gamma power is not universal, but dependent on the specific brain region, task epoch and nature of task performed. Many developmental EEG studies utilized resting state EEG that reflects the network at a different state compared to when it is actively engaged in a working memory task, or relied on passive sensory tasks, which revealed the greatest effects in lower cortical areas, including stronger adult 40-Hz auditory steady-state response dominantly generated in the auditory cortex and visually-evoked gamma (30-148 Hz) oscillation amplitude in the occipital lobe (Uhlhaas and Singer, 2011). Task-dependent changes involving higher-order cognitive functions often saw effects that were differentiable between frontal and parietal regions (Roux and Uhlhaas, 2014). The working memory load-dependent gamma power (50 – 100 Hz) change reported by Kornblith et al. (2016) differed between task epochs and brain regions. During and immediately after cue presentation, gamma power increased with higher memory load in the parietal cortex but decreased with higher memory load in the prefrontal cortex, while in the late delay period, there was no significant load-dependent gamma power changes in either brain areas. Uhlhaas et al. (2009) reported age-dependent gamma (30 – 75 Hz) power changes in a face perception task for parietal electrodes only. Furthermore, the task effect can be feature specific. Honkanen et al. (2015) saw load-dependent increases of gamma (40 – 72 Hz and 80 – 120 Hz) power for color and shape features but a decrease for location features. At least some preliminary evidence exists for decreases in EEG gamma power in the delay period of the Oculomotor Delayed Response task across adolescent development (McKeon et al., 2020).

Oscillatory discharges increase early in postnatal development in non-human primates, as inferred by intracellular recording experiments in slice preparations (Gonzalez-Burgos et al., 2015). It is possible therefore that gamma oscillations follow an inverted U curve during development. Gamma power has been also described in other animal models, e.g. revealing a monotonic postnatal increase in prefrontal gamma power in some rodent studies (Bitzenhofer et al., 2020). However, key differences have been discovered between animal models in the developmental profile of excitatory-inhibitory circuits, including the lack of NMDA receptors on adult frontal interneurons (Wang and Gao, 2009) as well as GABA synaptogenesis and functional maturation being complete well before the onset of adolescence in the rodents (Le Magueresse and Monyer, 2013).

Abnormally lower gamma power is commonly observed in schizophrenia (Woo et al., 2010; Uhlhaas and Singer, 2013), generally attributed to a shift of the excitation/inhibition balance towards a more excitable cortical state (Lisman, 2012). The decrease in gamma power is specific for stimulus presentation and task engagement in patients with schizophrenia (Uhlhaas and Singer, 2010); spontaneous gamma power may be elevated in these patients, as is during psychotic episodes and auditory hallucinations (Baldeweg et al., 1998; Spencer et al., 2009; Grent-’t-Jong et al., 2018). In this context, our results would suggest that the primate adolescence represents a state in the opposite end of the excitation/inhibition spectrum, dominated by greater inhibition, which is again consistent with the idea of elevated inhibitory drive in adolescence (Zhou et al., 2014). In turn, this result would suggest that schizophrenia represents an aberrant, excessive decrease in inhibition, rather than a prolonged adolescent-like state that failed to mature.

Through both empirical and modeling studies, a robust association has been established between higher gamma power and recurrent activity. Indeed, we observed stronger gamma power at recording sites with more neuronal information encoded. More subtly, the gamma power and neuronal firing of local circuits became more aligned in the adult. Both findings agree with the assumed role of gamma power at the microcircuit level. However, the overall direction of change in gamma power between stages was inverted against that of neuronal firing and behavior performance. These results suggest that during brain maturation, there might be systematic changes at the molecular and/or network level prominent enough to mask changes in features specific to a cognitive function of the brain region of interest. These effects would be especially challenging to differentiate when only macroscopic measures of neural activity are adopted. Bridging the processes underlying rhythmicity from the neuron to the circuit level will be an important goal of future studies.

## Acknowledgements

Research reported in this paper was supported by the National Institute of Mental Health of the National Institutes of Health under award numbers R01 MH117996 and R01 MH116675. We wish to thank Austin Lodish and Junda Zhu for technical and analytical help and André Bastos, Beatriz Luna, and David Blake for helpful comments on the manuscript.

## REFERENCES

Anderson SA, Classey JD, Conde F, Lund JS, Lewis DA (1995) Synchronous development of pyramidal neuron dendritic spines and parvalbumin-immunoreactive chandelier neuron axon terminals in layer III of monkey prefrontal cortex. Neuroscience 67:7–22.

Baldeweg T, Spence S, Hirsch SR, Gruzelier J (1998) Gamma-band electroencephalographic oscillations in a patient with somatic hallucinations. Lancet 352:620–621.

Bitzenhofer SH, Pöpplau JA, Hanganu-Opatz I (2020) Gamma activity accelerates during prefrontal development. eLife 9:e56795.

Bourgeois JP, Goldman-Rakic PS, Rakic P (1994) Synaptogenesis in the prefrontal cortex of rhesus monkeys. Cereb Cortex 4:78–96.

Bunge SA, Dudukovic NM, Thomason ME, Vaidya CJ, Gabrieli JD (2002) Immature frontal lobe contributions to cognitive control in children: evidence from fMRI. Neuron 33:301–311.

Burgund ED, Lugar HM, Miezin FM, Schlaggar BL, Petersen SE (2006) The development of sustained and transient neural activity. Neuroimage 29:812–821.

Buzsaki G, Wang XJ (2012) Mechanisms of gamma oscillations. Annu Rev Neurosci 35:203–225.

Compte A, Brunel N, Goldman-Rakic PS, Wang XJ (2000) Synaptic mechanisms and network dynamics underlying spatial working memory in a cortical network model. Cereb Cortex 10:910–923.

Compte A, Constantinidis C, Tegner J, Raghavachari S, Chafee MV, Goldman-Rakic PS, Wang XJ (2003) Temporally irregular mnemonic persistent activity in prefrontal neurons of monkeys during a delayed response task. J Neurophysiol 28:3441–3454.

Constantinidis C, Luna B (2019) Neural Substrates of Inhibitory Control Maturation in Adolescence. Trends Neurosci 42:604–616.

Davidson MC, Amso D, Anderson LC, Diamond A (2006) Development of cognitive control and executive functions from 4 to 13 years: evidence from manipulations of memory, inhibition, and task switching. Neuropsychologia 44:2037–2078.

Dienel SJ, Lewis DA (2019) Alterations in cortical interneurons and cognitive function in schizophrenia. Neurobiol Dis 131:104208.

Fries P (2009) Neuronal Gamma-Band Synchronization as a Fundamental Process in Cortical Computation. Annual Review of Neuroscience 32:209–224.

Fry AF, Hale S (2000) Relationships among processing speed, working memory, and fluid intelligence in children. Biol Psychol 54:1–34.

Gathercole SE, Pickering SJ, Ambridge B, Wearing H (2004) The structure of working memory from 4 to 15 years of age. Dev Psychol 40:177–190.

Giedd JN, Rapoport JL (2010) Structural MRI of Pediatric Brain Development: What Have We Learned and Where Are We Going? Neuron 67:728–734.

Goldman-Rakic PS (1994) Working memory dysfunction in schizophrenia. J Neuropsychiatry Clin Neurosci 6:348–357.

Gonzalez-Burgos G, Miyamae T, Pafundo DE, Yoshino H, Rotaru DC, Hoftman G, Datta D, Zhang Y, Hammond M, Sampson AR, Fish KN, Ermentrout GB, Lewis DA (2015) Functional Maturation of GABA Synapses During Postnatal Development of the Monkey Dorsolateral Prefrontal Cortex. Cereb Cortex 25:4076–4093.

Grent-’t-Jong T, Gross J, Goense J, Wibral M, Gajwani R, Gumley AI, Lawrie SM, Schwannauer M, Schultze-Lutter F, Navarro Schroder T, Koethe D, Leweke FM, Singer W, Uhlhaas PJ (2018) Resting-state gamma-band power alterations in schizophrenia reveal E/I-balance abnormalities across illness-stages. Elife 7.

Hoftman GD, Lewis DA (2011) Postnatal developmental trajectories of neural circuits in the primate prefrontal cortex: identifying sensitive periods for vulnerability to schizophrenia. Schizophr Bull 37:493–503.

Honkanen R, Rouhinen S, Wang SH, Palva JM, Palva S (2015) Gamma Oscillations Underlie the Maintenance of Feature-Specific Information and the Contents of Visual Working Memory. Cerebral Cortex 25:3788–3801.

Howard MW, Rizzuto DS, Caplan JB, Madsen JR, Lisman J, Aschenbrenner-Scheibe R, Schulze-Bonhage A, Kahana MJ (2003) Gamma Oscillations Correlate with Working Memory Load in Humans. Cereb Cortex 13:1369–1374.

Huttenlocher PR, Dabholkar AS (1997) Regional differences in synaptogenesis in human cerebral cortex. The Journal of comparative neurology 387.

Jensen O, Kaiser J, Lachaux JP (2007) Human gamma-frequency oscillations associated with attention and memory. Trends Neurosci 30:317–324.

Kajikawa Y, Schroeder Charles E (2011) How Local Is the Local Field Potential? Neuron 72:847–858.

Klingberg T, Forssberg H, Westerberg H (2002) Training of working memory in children with ADHD. J Clin Exp Neuropsychol 24:781–791.

Kornblith S, Buschman TJ, Miller E (2016) Stimulus Load and Oscillatory Activity in Higher Cortex. Cerebral cortex.

Kwon H, Reiss AL, Menon V (2002) Neural basis of protracted developmental changes in visuo-spatial working memory. Proc Natl Acad Sci U S A 99:13336–13341.

Le Magueresse C, Monyer H (2013) GABAergic interneurons shape the functional maturation of the cortex. Neuron 77:388–405.

Lewis DA, Cruz D, Eggan S, Erickson S (2004) Postnatal development of prefrontal inhibitory circuits and the pathophysiology of cognitive dysfunction in schizophrenia. Annals of the New York Academy of Sciences 1021:64–76.

Li S, Zhou X, Constantinidis C, Qi XL (2020) Plasticity of Persistent Activity and Its Constraints. Front Neural Circuits 14:15.

Lisman J (2012) Excitation, inhibition, local oscillations, or large-scale loops: what causes the symptoms of schizophrenia? Curr Opin Neurobiol 22:537–544.

Luna B, Thulborn KR, Munoz DP, Merriam EP, Garver KE, Minshew NJ, Keshavan MS, Genovese CR, Eddy WF, Sweeney JA (2001) Maturation of widely distributed brain function subserves cognitive development. Neuroimage 13:786–793.

Lundqvist M, Herman P, Warden MR, Brincat SL, Miller EK (2018) Gamma and beta bursts during working memory readout suggest roles in its volitional control. Nat Commun 9:394.

Lundqvist M, Rose J, Herman P, Brincat SL, Buschman TJ, Miller EK (2016) Gamma and Beta Bursts Underlie Working Memory. Neuron 90:152–164.

McKeon S, Calabro F, Luna B (2020) Development of EEG-derived spectral processing of working memory through adolescence In: Flux Congress, pp 82–82.

Olesen PJ, Nagy Z, Westerberg H, Klingberg T (2003) Combined analysis of DTI and fMRI data reveals a joint maturation of white and grey matter in a fronto-parietal network. Brain Res Cogn Brain Res 18:48–57.

Olesen PJ, Macoveanu J, Tegner J, Klingberg T (2007) Brain activity related to working memory and distraction in children and adults. Cereb Cortex 17:1047–1054.

Pesaran B, Pezaris JS, Sahani M, Mitra PP, Andersen RA (2002) Temporal structure in neuronal activity during working memory in macaque parietal cortex. Nat Neurosci 5:805–811.

Ray S, Maunsell JH (2011) Different origins of gamma rhythm and high-gamma activity in macaque visual cortex. PLoS Biol 9:e1000610.

Roux F, Uhlhaas PJ (2014) Working memory and neural oscillations: alpha-gamma versus theta-gamma codes for distinct WM information? Trends Cogn Sci 18:16–25.

Spencer KM, Niznikiewicz MA, Nestor PG, Shenton ME, McCarley RW (2009) Left auditory cortex gamma synchronization and auditory hallucination symptoms in schizophrenia. BMC Neurosci 10:85.

Uhlhaas PJ, Singer W (2010) Abnormal neural oscillations and synchrony in schizophrenia. Nat Rev Neurosci 11:100–113.

Uhlhaas PJ, Singer W (2011) The Development of Neural Synchrony and Large-Scale Cortical Networks During Adolescence: Relevance for the Pathophysiology of Schizophrenia and Neurodevelopmental Hypothesis. Schizophrenia Bulletin 37:514–523.

Uhlhaas PJ, Singer W (2013) High-frequency oscillations and the neurobiology of schizophrenia. Dialogues Clin Neurosci 15:301–313.

Uhlhaas PJ, Roux F, Rodriguez E, Rotarska-Jagiela A, Singer W (2010) Neural synchrony and the development of cortical networks. Trends Cogn Sci 14:72–80.

Uhlhaas PJ, Roux F, Singer W, Haenschel C, Sireteanu R, Rodriguez E (2009) The development of neural synchrony reflects late maturation and restructuring of functional networks in humans. Proc Natl Acad Sci U S A 106:9866–9871.

Ullman H, Almeida R, Klingberg T (2014) Structural maturation and brain activity predict future working memory capacity during childhood development. J Neurosci 34:1592–1598.

Wang H-X, Gao W-J (2009) Cell Type-Specific Development of NMDA Receptors in the Interneurons of Rat Prefrontal Cortex. Neuropsychopharmacology 34:2028–2040.

Wang X-J (2010) Neurophysiological and Computational Principles of Cortical Rhythms in Cognition. Physiological Reviews 90:1195–1268.

Woo T-UW, Spencer K, McCarley RW (2010) Gamma Oscillation Deficits and the Onset and Early Progression of Schizophrenia. Harvard Review of Psychiatry 18:173–189.

Zhou X, Qi XL, Constantinidis C (2016a) Distinct Roles of the Prefrontal and Posterior Parietal Cortices in Response Inhibition. Cell Rep 14:2765–2773.

Zhou X, Zhu D, Qi XL, Lees CJ, Bennett AJ, Salinas E, Stanford TR, Constantinidis C (2013) Working Memory Performance and Neural Activity in the Prefrontal Cortex of Peri-pubertal Monkeys. J Neurophysiol 110:2648–2660.

Zhou X, Zhu D, King SG, Lees CJ, Bennett AJ, Salinas E, Stanford TR, Constantinidis C (2016b) Behavioral response inhibition and maturation of goal representation in prefrontal cortex after puberty. Proc Natl Acad Sci U S A 113:3353–3358.

Zhou X, Zhu D, Qi XL, Li S, King SG, Salinas E, Stanford TR, Constantinidis C (2016c) Neural correlates of working memory development in adolescent primates. Nat Commun 7:13423.

Zhou X, Zhu D, Katsuki F, Qi XL, Lees CJ, Bennett AJ, Salinas E, Stanford TR, Constantinidis C (2014) Age-dependent changes in prefrontal intrinsic connectivity. Proc Natl Acad Sci U S A 111:3853–3858.

